# Twitches emerge postnatally during quiet sleep in human infants and are synchronized with sleep spindles

**DOI:** 10.1101/2021.02.17.431688

**Authors:** Greta Sokoloff, James C. Dooley, Ryan M. Glanz, Rebecca Y. Wen, Meredith M. Hickerson, Laura G. Evans, Haley M. Laughlin, Keith S. Apfelbaum, Mark S. Blumberg

**Author notes:** Corresponding authors: Greta Sokoloff, Ph.D., Mark S. Blumberg, Ph.D. Lead Contact: Greta Sokoloff.

## Abstract

In humans and other mammals, the stillness of sleep is punctuated by bursts of rapid eye movements (REMs) and myoclonic twitches of the limbs [1]. Contrary to the notion that twitches are mere by-products of dreams, sensory feedback arising from twitching limbs provides a rich and unique source of activation to the developing sensorimotor system [2]. In fact, it is partly because of the behavioral activation of REM sleep that this state is also called active sleep (AS), in contrast with the behavioral quiescence that gives quiet sleep (QS)—the second major stage of sleep—its name. In human infants, for which AS occupies eight or more hours of each day [3], limb twitching is one among several components that help to identify the state [4-7]; nonetheless, we know relatively little about the structure and functions of twitching across development. Recently, in sleeping infants over the first seven postnatal months [8], we documented a pronounced shift in the temporal expression of twitching beginning around three months of age that suggested a qualitative shift in how twitches are produced. Here, we combine behavioral assessments of twitching with high-density electroencephalography (EEG) and demonstrate that this shift reflects the developmental emergence of limb twitches during QS. Twitches during QS are not only unaccompanied by REMs, but they also occur synchronously with sleep spindles, a hallmark of QS. As QS-related twitching increases with age, sleep spindle rate also increases along the sensorimotor strip. The emerging synchrony between subcortically generated twitches and cortical oscillations suggests the development of functional connectivity among distant sensorimotor structures, with potential implications for detecting and explaining atypical developmental trajectories.

## Results and Discussion

We recorded infants’ sleep during the daytime in the laboratory. In all sessions, infants slept in a supine, semi-reclined position, as described previously [8]. Infants were fitted with a high-density electrode cap for measurement of EEG and movements were recorded using a video camera from a frontal view. A total of 22 infants (15 boys, 7 girls) between one week and seven months of age participated and slept during the session, yielding a total of 55 sessions (see Table S1 for descriptive data).

Representative 18-minute periods of sleep and wake are presented in Figure 1 for infants at three ages. At one month (Figure 1A), the high density of REMs and limb and facial twitches clearly demarcate the state of AS. QS is identified by the near-total absence of REMs and twitches in an infant that otherwise exhibits other behavioral manifestations of sleep; there is also a small increase in power in the low-frequency EEG domain (delta: 0.5-4 Hz). At three (Figure 1B) and six (Figure 1C) months of age, sleep spindles—brief thalamocortical oscillations at 12-14 Hz—occur approximately once every 10 seconds [9]. Sleep spindles and increased delta power become defining features of QS at these older ages [4].

**Figure 1.**
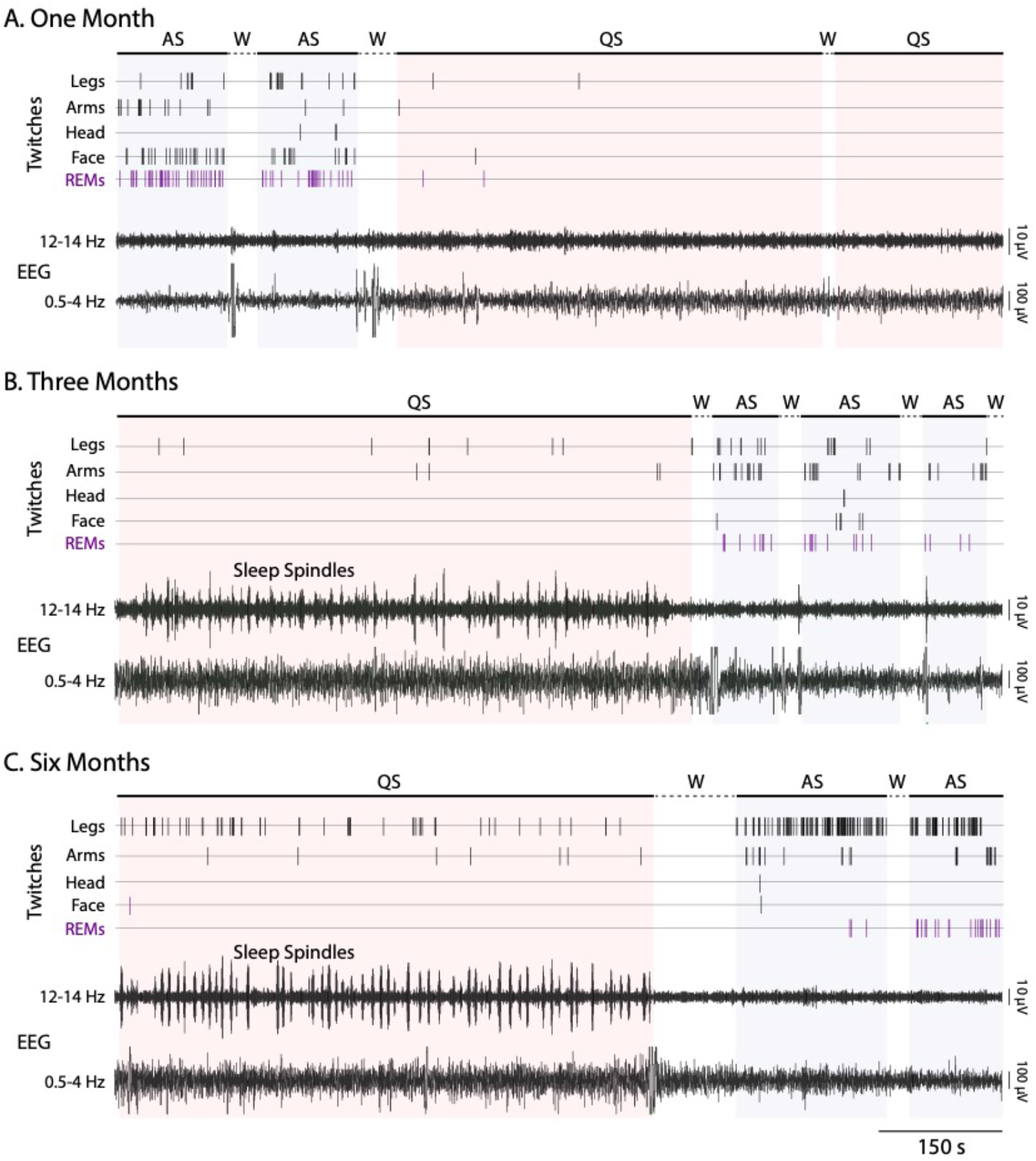
Representative records of behavior and EEG activity in sleeping human infants. (A) Continuous ∼18-minute record of active sleep (AS; blue shading), quiet sleep (QS; red shading), and wake (W) for a one-month-old infant. Twitches of the legs, arms, head, and face are shown (black ticks), as are rapid eye movements (REMs; purple ticks). EEG records, recorded from the C3 electrode site, are filtered for spindle (12-14 Hz) and delta (0.5-4 Hz) frequencies. (B) Same as in (A) except for a three-month-old infant. (C) Same as in (A) except for a six-month-old infant.

As expected, all bouts of AS were associated with high-density limb twitching and REMs. However, we also observed the developmental emergence of limb twitches during sleep-spindle-rich periods of QS (Figure 1B and C; Movie S1). Upon close visual inspection, experimenters were unable to distinguish between twitches during AS and QS. This observation was initially difficult to reconcile with the conventional, defining association between twitches and AS in humans, rats, and other infant mammals [4,5,10]. One approach to reconciling the temporal overlap of an AS component (e.g., twitches) with a QS component (e.g., sleep spindles) is to label the period of sleep as “indeterminate” or “transitional” [4]. Rather than adopting this conventional approach, however, we instead assessed whether twitches during QS exhibit developmental or spatiotemporal features that distinguishing them from twitches during AS.

We first compared rates of twitching during QS and AS for each sleep session (Figure 2A). Whereas AS-related twitches occurred at a high and stable rate across all ages, the rate of QS-related twitches only began to increase after three months of age. A mixed-effects model revealed a significant main effect of sleep state (B=8.28, SE=1.03, t(55.6)=8.04, p<.0001) but not age (B=0.83, SE=0.43, t(21.5)=1.93, p<0.07). The interaction was significant (B=-1.55, SE=0.61, t(57.6)=-2.55, p=.013) and arose because of a larger effect of age during QS than during AS. As expected, REMs occurred at a high rate during AS and occurred rarely during QS at all ages (Figure 2B). Although a few studies have described twitching during QS in human adults [11-13], the rates reported in those studies appear much lower than those observed here in infants.

**Figure 2.**
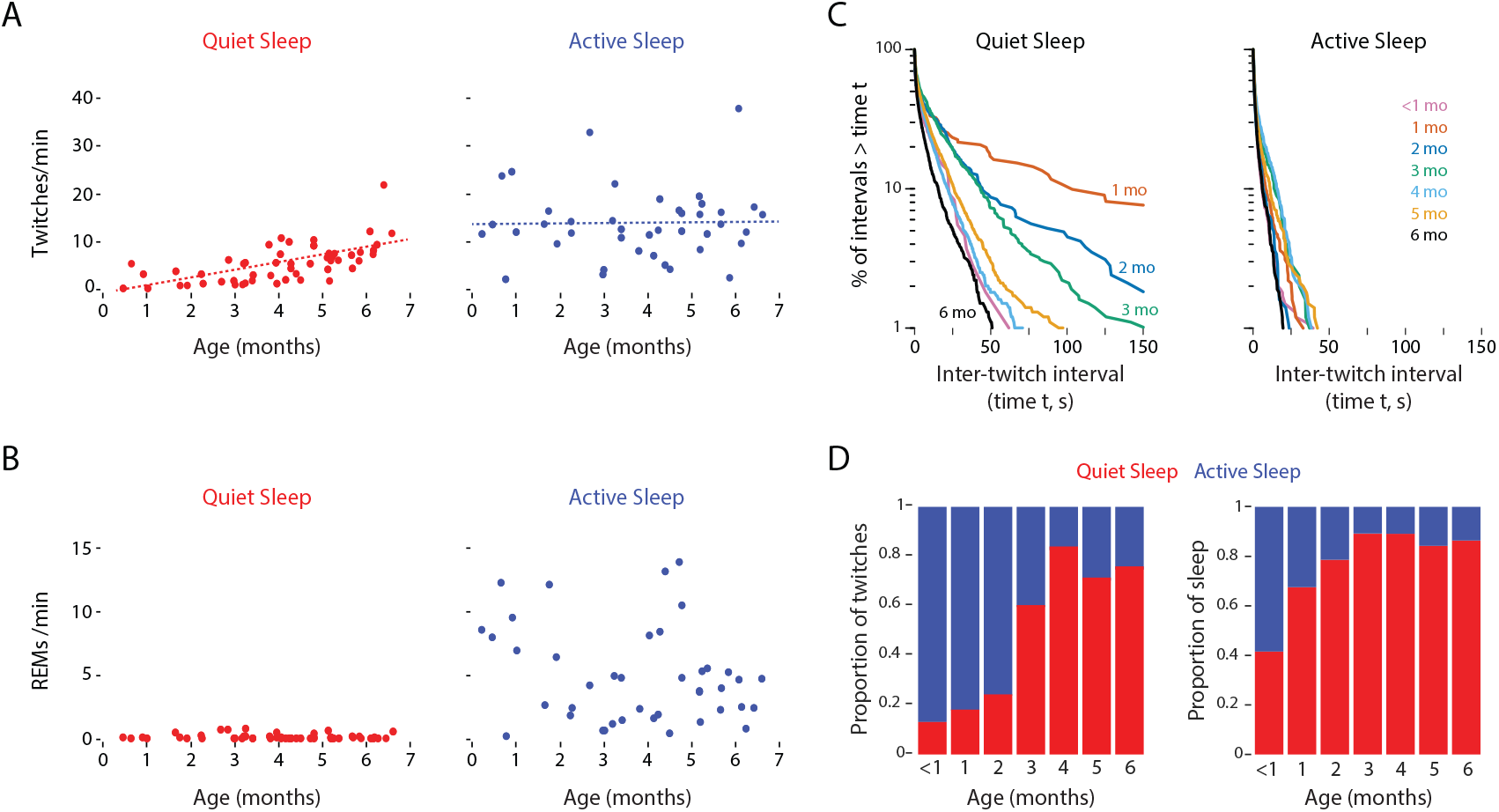
Temporal organization of QS- and AS-related twitching over development. (A) Scatterplots showing the rate of twitching during QS (red) and AS (blue) for each sleep session. The dotted lines show the model-predicted estimates for population means of twitch rate by age and sleep state. (B) Scatterplots showing the rate of rapid eye movements (REMs) during QS (red) and AS (blue) for each sleep session (C) Log-survivor plots of inter-twitch intervals (ITIs) during periods of QS and AS within each age group (see Figure S1B for sample sizes for each plot). ITIs were pooled across sleep sessions within each month of age. (D) Left: Proportion of twitches during QS (red) and AS (blue) within each age group for data pooled across sleep sessions. Right: Proportion of sleep time in QS (red) and AS (blue) within each age group for data pooled across sleep sessions.

The representative data in Figure 1 suggest that twitches during AS are more likely to occur in rapid succession than twitches during QS. Thus, to assess differences in the temporal structure of twitching during QS and AS, we generated log-survivor plots of inter-twitch intervals (ITIs) during QS and AS across age. Linearity on such semi-log plots denotes a constant average rate of twitching, with steeper slopes indicating higher rates. First, all these plots have their steepest slopes at the shortest ITIs, indicative of the fact that most twitches tend to occur in rapid succession; indeed, at all ages for both QS and AS, the majority of ITIs were less than 1 s. Second, consistent with the data in Figure 1, the ITI survivor plots for AS—across all age groups—exhibited slopes that were steeper than those for twitching during QS, thus indicating that twitches during AS occurred at higher average rates (Figures 2C and S1A-B). Between one and four months of age, the slopes for the QS distributions gradually approached those for the AS distributions.

Consistent with the foregoing analyses, ≤20% of all twitches in our sample occurred during QS through two months of age before increasing to nearly 80% by four months of age (Figure 2D, left). Moreover, QS-related twitching began to increase between two and three months of age as the proportion of sleep time in QS plateaued at ∼80% of total sleep time (Figure 2D, right).

All together, these findings indicate that QS-related twitches emerge postnatally and exhibit temporal features across age that distinguish them from AS-related twitches. However, when we performed a more focused assessment of the distribution of twitches across the body in the five-month-old infants, there were no differences between QS and AS: For both states, twitches were most likely to occur in fingers and toes, followed by hands and feet, shoulders/elbows and hips/knees, and face and head (Figure S1C). This result suggests that twitches during QS and AS are produced by the same or overlapping neural mechanisms.

Given that twitches and sleep spindles co-occur during QS and that both increase in frequency over these postnatal ages (see Figure 1), we next quantified sleep spindles (Figure S2A and Table S1) and assessed their temporal association with twitches. As described previously in human infants [9,14], sleep spindles occurred with peak frequencies of 12-14 Hz (Figure 3A). The rate of sleep spindles increased between one and four months of age (Figure 3B); survivor analysis of inter-spindle intervals revealed a development shift in the production of sleep spindles between one and three months of age, after which the distributions stabilized at high constant average rates (Figure 3C); also, as sleep spindles emerged, they exhibited a dominant periodicity of ∼10 s (Figure S2B), as described previously [9]. Visual inspection (Figure 3D) and quantitative analysis (Figure 3E) of spindle power indicated focal increases, beginning around three months of age, at electrode sites that lie along the sensorimotor strip (Figure 3D). A mixed-effects model revealed significant main effects of sleep state (B=-3.82, SE=0.43, t(72.3)=-8.83, p<.0001) and age (B=0.99, SE=0.17, t(26.4)=5.92, p<0.0001), and a significant interaction (B=-0.73, SE=0.25, t(73.6)=-2.86, p=.0055); the interaction arose because of a larger effect of age during QS than during AS. A parallel analysis of delta power during QS (0.5-4 Hz) revealed significant main effects of age and sleep state, but not a significant interaction (see Figure S3 for detailed results).

We assessed each bout of QS for all sessions and sorted them from shortest (<1 min) to longest (40 min), noting the occurrence of every limb twitch and sleep spindle (Figure S4). Within each QS bout, whereas sleep spindles distributed evenly across the bout, twitches occurred more sporadically. By four months of age, twitches tended to occur at higher rates at the beginning of each bout. Delta power exhibited an inverse relationship with twitching, tending to begin each bout at lower values and increase thereafter.

**Figure 3.**
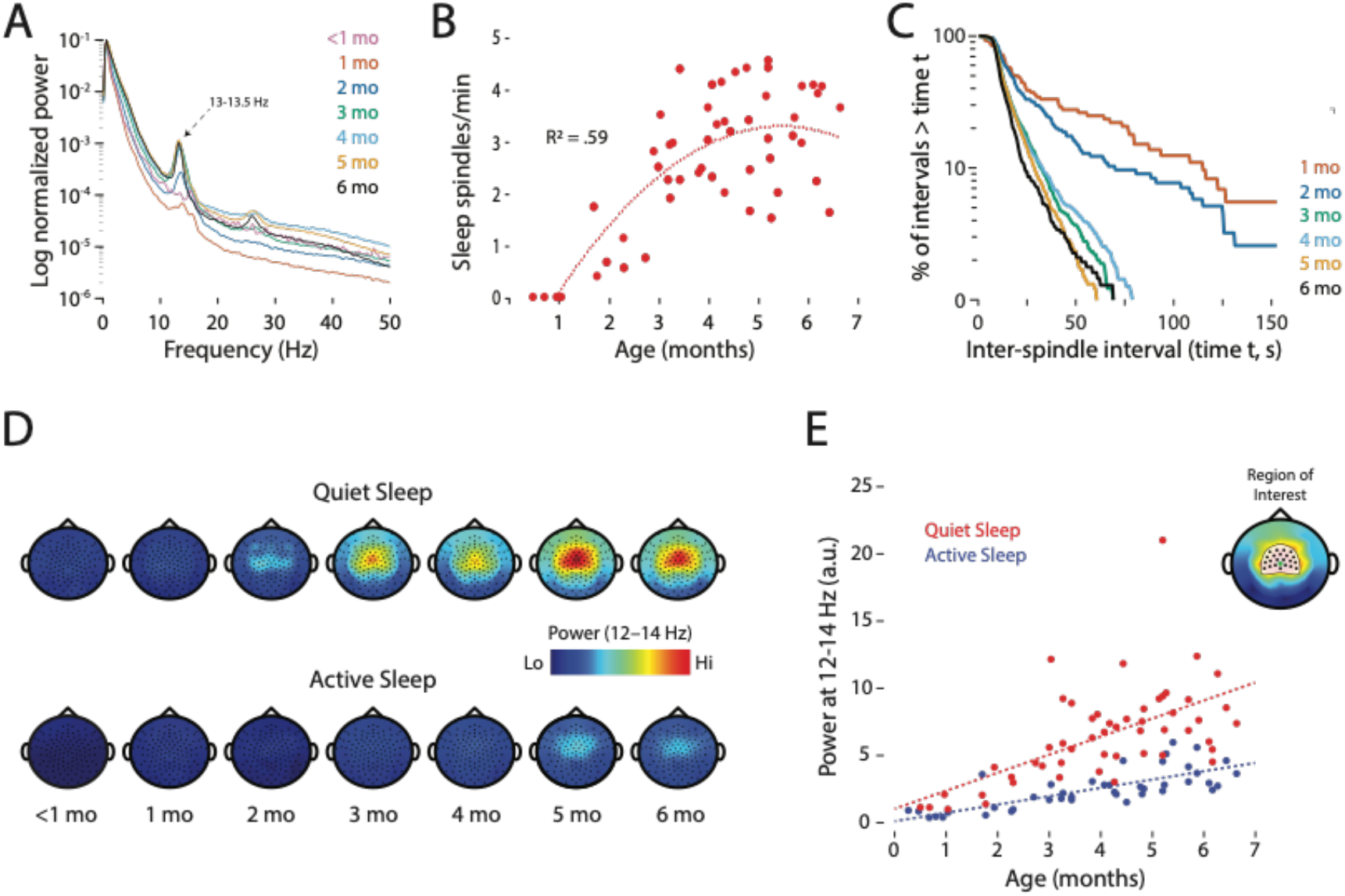
Increase in the rate and power of sleep spindles over development. (A) Power spectra at representative electrode sites in centro-parietal cortex (C3 or C4), averaged across sleep sessions at each age. Note the frequency peaks at 13-13.5 Hz, particularly at 2-6 months of age. (B) Scatterplot showing the rate of sleep spindles during QS for each sleep session. Best-fit polynomial line and R^2^ value is shown. (C) Log-survivor plots of inter-spindle intervals during periods of QS in infants from 1 to 6 months of age. Intervals were pooled across all sleep sessions at each age (Ns = 72-1189 inter-spindle intervals). (D) Topoplots showing sleep spindle power (12-14 Hz) during QS and AS. Data averaged across sleep sessions at each age. (E) Power at sleep-spindle frequency (12-14 Hz; arbitrary units) averaged within a region of interest (inset; Cz electrode indicated by a green dot) during QS (red) and AS (blue) for each sleep session. The dotted lines show the model-predicted estimates for population means of spindle power by age and sleep state.

In infant and adult animals, sleep spindles are implicated in learning and plasticity, including plasticity in the sensorimotor system [15]. Accordingly, a close temporal association between spindles and twitches could suggest that they function together to promote sensorimotor development. Thus, we next determined whether spindles and twitches are temporally associated; in the periods surrounding twitches of the arms or legs, we calculated the likelihood of a sleep spindle at somatotopically relevant electrode sites (Figure 4A-B). By three months of age, peak probabilities of sleep spindles coincided with the occurrence of arm or leg twitches (Figure 4C). Next, we compared the observed probabilities of detecting a sleep spindle within 3 s of a twitch and compared those probabilities to expected values based on shuffled data (Figure 4D). Paired t tests revealed significant differences between expected and observed values for arms and legs from three to six months of age, but not at two months of age. Thus, as QS-related twitches emerge, they become temporally coupled with sleep spindles.

**Figure 4.**
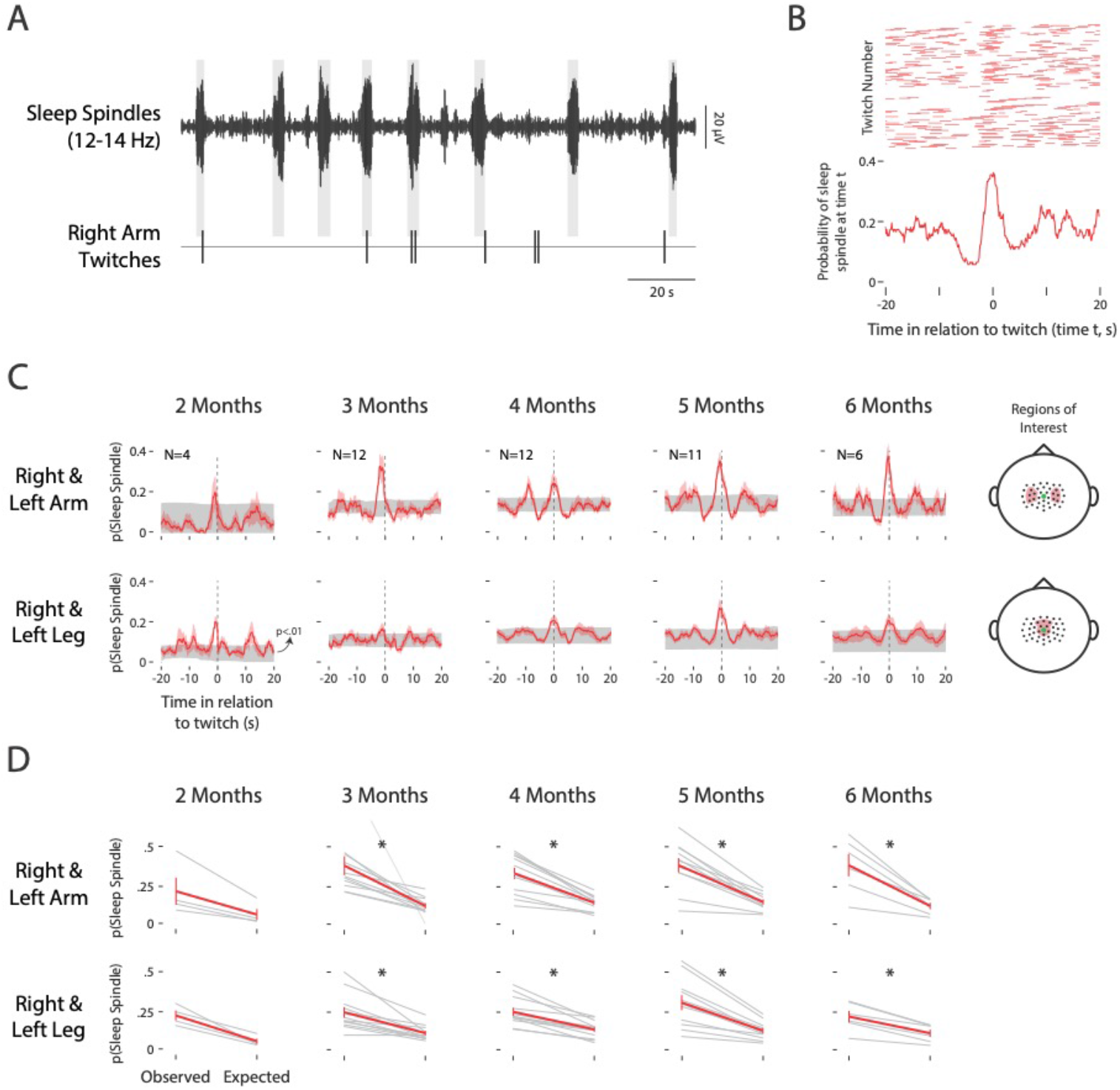
Sleep spindles occur in synchrony with arm and leg twitches.(A) Representative 155-s record from a 6-month-old infant to show twitches of the right arm in relation to spindles recorded from C3. Shaded rectangles indicate the duration of the spindles. (B) Top: Raster plot showing the occurrence of sleep spindles recorded from C4 over a +20-s window surrounding twitches of the left arm during one sleep session for a 6-month-old infant; each of the 138 rows denotes data for a different twitch. Bottom: The probability of detecting a sleep spindle over a +20-s peri-twitch window for all 138 twitches. (C) Mean probability (red lines) of detecting a sleep spindle plotted against time in relation to a twitch of the right or left arm (top row) and right or left leg (bottom row) at two through six months of age. The standard errors around the means are shown as red shading. For each plot, the 99% confidence interval is shown in gray (p<.01). Also shown is the number (N) of sleep sessions for each age group that were included in these analyses. Far right: Electrode selections for the analyses in each row are indicated by red shading. The electrode selection was in the contralateral hemisphere for right or left arm twitches, and in the same medial location for both right and left leg twitches. The Cz electrode is indicated by a green dot. (D) For each sleep session, we calculated the maximum probability of observing a sleep spindle within +3 s of a twitch of the right and left arms (top row) and right and left legs (bottom row). Expected values were determined for each sleep session by shuffling the data 1000 times. Individual session data (gray lines) and mean data (red lines; with standard errors) are shown. * significant difference between observed and expected (Arms: two months: t_3_= 2.86, NS; three months: t_10_= 8.84, p < .005; four months: t_11_= 7.29, p < .005; five months: t_10_= 6.73, p < .005; six months: t_5_= 4.84, p < .005; Legs: two months: t_3_= 5.15, NS; three months: t_10_= 4.32, p < .005; four months: t_11_= 6.33, p < .005; five months: t_10_= 5,72, p < .005; six months: t_5_= 6.15, p < .005).

As expected, during AS we did not observe any twitch-associated spindle activity in these subjects. However, AS-related EEG activity does occur in premature infants and newborns [16,17]; at those earlier ages, activity takes the form of “spindle bursts,” brief thalamocortical oscillations that resemble sleep spindles in some ways, including their dominant frequency [18]. As studied most extensively in infant rats, AS-related twitches are produced by motor structures in the brainstem and trigger sensory feedback that cascades throughout the brain, from medulla to sensorimotor cortex [2]. As in human infants, spindle bursts in rats disappear postnatally, apparently due to the onset of inhibitory mechanisms in the thalamus [19]. It is not until after AS-related spindle bursts are no longer detected that QS-related sleep spindles emerge and, as shown here, are accompanied by the onset of QS-related twitching.

Whereas spindle bursts in sensorimotor cortex are clearly sensory responses to movement, this does not appear to be the case for sleep spindles. Nor have sleep spindles been implicated in the production of movement. Thus, it is possible that the twitches observed here in QS and AS are produced by similar brainstem mechanisms, with those mechanisms being differentially modulated in a state-dependent manner across age. If so, then the temporal association between twitches and sleep spindles (Figure 4) could reflect emerging communication among brainstem and cortical motor structures, including primary motor cortex (M1). In fact, in adult rats, sleep spindles increase the likelihood that neurons in M1 fire synchronously [20] and contribute to M1 activity associated with the learning of a skilled motor task [21].

In mammals, M1 develops much later than is commonly realized [22,23]. In humans, the late onset of M1 functionality dramatically reveals itself in infants who have experienced a perinatal stroke, which is the leading cause of hemiplegic cerebral palsy (CP) [24]. Because of the protracted postnatal development of M1 and its descending connections, the disabling effects of CP are not evident until at least 4–8 months after birth [25]. At 2-4 months of age, before the onset of CP symptoms, the wake-related “fidgety” movements that are typically observed are curiously absent [26]. Accordingly, we propose that our two key findings—the postnatal emergence of QS-related twitches and their close temporal association with sleep spindles in sensorimotor cortex—reflect a cortical motor system that is just beginning to establish functional connectivity with subcortical motor structures. Accordingly, QS-related twitching has the potential to provide additional leverage for detecting and understanding typical and atypical motor development.

## Acknowledgments

We thank Sampsa Vanhatalo and Karen Adolph for helpful comments. Research supported by NIH R21-HD095153 and R37-HD081168.

## Author Contributions

Conceptualization, G.S. and M.S.B.; Methodology, G.S. and M.S.B.; Formal Analysis, G.S., J.C.D., R.M.G., K.S.A., and M.S.B.; Investigation, G.S., R.Y.W., M.M.H., L.G.E., and H.M.L.; Resources, G.S. and M.S.B.; Data Curation, G.S. and R.Y.W.; Writing - Original Draft, G.S., R.Y.W., J.C.D, R.M.G., K.S.A., and M.S.B.; Writing—Reviewing and Editing, G.S., J.C.D., R.M.G., R.Y.W., M.M.H., K.S.A., and M.S.B.; Visualization, G.S., J.C.D., R.M.G., and M.S.B.; Supervision, G.S. and M.S.B.; Project Administration, G.S. and M.S.B.; Funding Acquisition, G.S. and M.S.B.

## STAR METHODS

### CONTACT FOR REAGENT AND RESOURCE SHARING

Further information and requests for resources and reagents should be directed to, and will be fulfilled by, the lead contact, Dr. Greta Sokoloff.

### SUBJECT DETAILS

A total of 22 infants (15 boys, 7 girls) between one week and seven months of age participated, yielding a total of 58 sleep sessions; eight infants contributed one session and the remaining 16 infants contributed 2-6 sessions. Two sessions were excluded from analysis because the infants slept for fewer than five min and one session was excluded for technical problems with the EEG signal. All but two infants were born at term. All study procedures were approved by the Institutional Review Board at the University of Iowa.

## METHOD DETAILS

### Study Procedure

We recorded infants’ sleep and EEG activity during the daytime in the laboratory. Infants arrived at the laboratory with a caregiver. Upon arrival, infants were changed into a plain black onesie and the experimenters measured their head circumference and visualized the vertex by determining the intersection of the nasion and inion. An EEG sensor cap (Electrical Geodesics, Inc., Eugene, OR), sized to the infant’s head and presoaked in a warm potassium chloride and baby shampoo solution, was fitted in place. Infants were then placed in a supine, semi-reclined position in a baby lounger and allowed to go to sleep. While their infants slept, caregivers sat in the testing room near their infants or in a nearby waiting room (equipped with a baby video monitor) and completed a demographics questionnaire.

### EEG and Video Data Acquisition

Cortical EEG was recorded continuously using a 128-channel HydroCel geodesic sensor net connected to a Net Amps 200 Amplifier (Electrical Geodesics, Inc.). Data were acquired at 1000 samples/s to a computer running Net Station software. When infants fell asleep, electrode impedances were checked to ensure that the majority of values were below 100 KΩ. For most sessions, impedances were also checked at the end of the recording session. EEG recordings were referenced online to Cz and later re-referenced depending on the analysis. A high-definition video camera (Model Q6055, Axis Communications, Lund, Sweden), placed to provide a frontal view of the infants, recorded movements of the eyes, face, head, and limbs in synchrony with the EEG data; video data were recorded at 30 frames/s. Recordings continued for as long as infants remained asleep or, when aroused, returned to sleep quickly. Finally, for a subset of infants, respiratory data were also acquired using a piezoelectric sensor (Pico Movement Sensor, Unimed Electrode Supplies Ltd, UK) attached to the infants’ outer garment at chest level.

### Behavioral Coding and Reliability

Twitches were coded using methods developed and described previously [8]. Two coders independently coded 100% of the video records for twitches. First, wake periods, startles, and arousals were identified for each session and used to segregate periods twitching. Next, coders scored the data in multiple passes to identify twitches of the face, head, left and right arms (including hands and fingers), and left and right legs (including feet and toes). Eye movements were also coded and designated as REMs when they had durations less than 1 s. Coders did not score movements during sleep that were produced incidentally due to respiration or movements of other limbs. Finally, coders were always blind to sleep state when they were scoring the occurrence of twitches.

Inter-observer reliability was assessed as described previously [8]. Briefly, we determined the time of occurrence of twitches detected by the two coders across all movement categories, applying a 0.5 s duration to each twitch. When comparing coders’ initial scores, Cohen’s kappa ranged from 0.321 to 0.973 (median: 0.6345). When coders disagreed, they independently reviewed the videos around the periods of interest and, when necessary, rescored the video. After this second pass through the data, Cohen’s kappa increased to 0.681-0.993 (median: 0.8635), indicating >80% agreement between the two coders. Finally, coders jointly made a final pass through the data record and all remaining discrepancies were resolved by mutual agreement.

For a subset of infants at five months of age (n=8), we reanalyzed the behaviorally scored data to determine the proportion of twitches during QS and AS that occurred in individual body segments. Thus, in addition to face and head twitches, we divided arms into shoulders/elbows, hands, and fingers, and legs into hips/knees, feet, and toes. For this analysis, we included only those infants that exhibited twitching during both QS and AS periods.

### EEG Data Acquisition

EEG data were band-pass filtered (0.5-60 Hz) and a 60-Hz notch was applied. Channels were re-referenced to the mean mastoid signal. EEG data were then imported into Spike2 (Cambridge Electronic Design, Ltd., Cambridge, UK) for sleep state determination. For all other analyses, raw, referenced EEG data were imported into MATLAB (The MathWorks, Inc., Natick, MA). A subset of 26 channels around the neck and ears were excluded from further analysis as these channels frequently contained movement artifact. EEG channels were also excluded when impedances were over 100 MΩ. Data for missing sites were interpolated when necessary for analysis.

### Sleep State Determination

Sleep was divided into periods of quiet sleep (QS) and active sleep (AS). Criteria for AS included the presence of rapid eye movements (REMs), low EEG amplitude, and the absence of sleep spindles. Criteria for QS included the presence of sleep spindles, high EEG amplitude, and the absence of REMs. Two of the three criteria had to be met to achieve a sleep state determination. Limb and face twitches were not included in these criteria.

For each sleep session during periods of REMs, an EEG amplitude threshold was established such that, when that threshold was exceeded, the EEG amplitude was considered “high.” After a transition from AS or wake, the first detected sleep spindle defined the onset of a QS bout. For periods of sleep when infants transitioned directly from AS to QS, the midpoint between the last twitch of the AS period and the first sleep spindle marked the state transition; conversely, for transitions from QS to AS, the midpoint between the last sleep spindle of the QS period and the first twitch of the AS period marked the state transition. The vast majority of sleep state transitions occurred after brief awakenings.

## QUANIFICATION AND STATISTICAL ANALYSIS

### Rates of Twitching and REMs

Counts for twitches (of the face, head, arms, and legs) and REMs were summed across periods of QS and AS and divided by the total time spent in each sleep state to obtain a measure of twitch and REM rates. Because REMs were a criterion for state determination, rate of REMs was not compared statistically between QS and AS.

Data for which age was treated as a continuous variable (i.e., rate of twitching, spindle power, delta power) were analyzed using linear mixed-effects models, using the lme4 package in R and the lmerTest package to estimate significance levels. For these analyses, sleep state was contrast coded (QS = -.5; AS = .5). Age was centered on 0 and included as a continuous factor. These variables served as independent variables for each model. Random intercepts included infant participant and, where possible, session number. In some instances, models that included session number as a random intercept produced singular fits, suggesting that the model was over-specified for the amount of data available; in such instances, only participant was included. In all models, more complex models that included random slopes produced singular fits or failed to converge.

The model used for testing age-related changes in the rate of twitching included sleep state and age as independent variables, and participant and sleep session as random intercepts. The full model structure is presented in Equation 1:

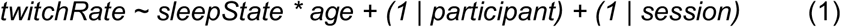

### Inter-twitch Intervals

For twitches during QS and AS within a sleep session, inter-twitch intervals (ITIs) were calculated as the time between successive twitches. ITIs were not included if there was an intervening startle, arousal, or at transitions between QS and AS. Twitches of all body segments (i.e., face, head, arms, and legs) were included in this analysis; REMs were not included. ITI data were pooled within age groups to generate log-survivor plots.

For the within-age comparisons between QS and AS, statistical differences in the survivor distributions were tested using the Mantel-Cox log-rank test in SPSS (IBM, Armonk, NY). Alpha was set at .05 and a Bonferroni procedure was used to correct for multiple comparisons.

### Spectral Analysis

To generate power spectra, QS periods were first identified based on the presence of sleep spindles in Spike2. Then, for a single site within the sensorimotor strip (e.g., electrodes 36 or 104 in the EGI sensor net system, corresponding to C3 or C4, respectively, in the 10-20 system), power spectra from 0.5 to 50 Hz were calculated using a 4-s Hanning window for the fast Fourier transform. Spectra were then averaged across sleep sessions within each age group.

### Spindle Detection and Analysis

Sleep spindles were detected at all EEG channels by first band-pass filtering the signals at 12-14 Hz (Figure S2A). Using a Hilbert transform of the filtered data, the amplitude of each signal was generated and periods of noise during QS were removed. The onset of a sleep spindle was marked when signal amplitude passed a threshold of 2x the median amplitude for at least 1 s. After this 1-s minimum duration, the spindle was terminated as soon as the signal decreased below the threshold, so long as it remained below the threshold for at least 1 s.

For each sleep session, the rate of sleep spindles was calculated by dividing the total number of sleep spindles at a single electrode site (electrodes 36 or 104) by the total number of minutes in QS. Log-survivor plots of inter-spindle intervals were created in the same way as were those for ITIs.

### Sleep Spindle Power and Delta Power

EEG data were first band-pass filtered for sleep spindles (12-14 Hz) and delta (0.5-4 Hz). To determine the distribution of spindle and delta power across the sensor net, the absolute value of the Hilbert-transformed EEG signal was calculated. Next, for each sleep session, median values during AS and QS at each site were calculated and topoplots were created. When data were missing or were outliers at individual sites, data were interpolated from nearby sites to produce a clean topoplot for each sleep session. Topoplots were then averaged (mean) across sleep sessions within each age group. To produce a single value for sleep-spindle and delta power for each sleep session, we calculated the median value over electrode sites within regions of interest (sleep spindles: electrodes 5, 6, 7, 12, 13, 20, 29, 30, 31, 36, 37, 42, 55, 80, 87, 93, 104, 105, 106, 111, 112, 118; delta: anterior electrodes 3, 4, 5, 6, 7, 10, 11, 12, 13, 16, 18, 19, 20, 23, 24, 28, 29, 30, 31, 36, 37, 55, 80, 87, 93, 104, 105, 106, 111, 112, 117, 118, 124; posterior electrodes 70, 74, 75, 81, 82, 83).

The age-related changes in sleep-spindle and delta power were analyzed using linear mixed-effects models, similar to the analysis of the rate of twitching described above. For spindle power, sleep state and age were used as independent variables and infant participant and session number were used as random intercepts. The full model structure is presented in Equation 2:

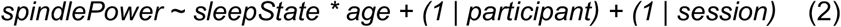

For delta power, sleep state and age were independent variables, and infant participant was used as a random intercept (including session number as a random intercept led to a singular model fit). The full model structure is presented in Equation 3:

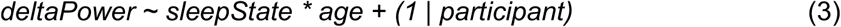

### Probability of Sleep Spindles in Relation to Twitching

We determined the probability that a sleep spindle occurred in the period before or after a twitch during all QS periods in infants at two months of age and older (only one of the three one-month-olds with sleep spindles exhibited sufficient sleep spindles and twitches for this analysis, so this age group was not included here). First, for right and left arm twitches, a region of interest was selected in the left and right somatosensory cortex, respectively (left: electrodes 29, 30, 36, 37, 42; right: electrodes 87, 93, 104, 105, 111); in contrast, for right and left leg twitches, a region of interest, located medially in somatosensory cortex, was selected (electrodes 6, 7, 13, 31, 55, 80, 106, 112). The regions of interest were selected based on visual inspection of the topoplots in relation to published descriptions of somatotopically relevant areas in the infant somatosensory cortex [27].

For each individual electrode site during periods of QS, we calculated the probability that a sleep spindle occurred at time t over a 40-s window (i.e., ±20 s around a twitch) in relation to twitches of the right and left arms and legs. For each limb, mean probabilities were then averaged across electrode sites within each region of interest. For the arms in each sleep session, to compensate for uneven numbers of right or left arm twitches, a weighted average was calculated to produce a single probability. Finally, mean probabilities for arms and legs were calculated across all sleep sessions within an age group.

For each probability plot, we calculated 99% confidence intervals (p<.01) using a Monte Carlo procedure. The timing of sleep spindles within for each channel was shuffled 1000 times using a procedure that retained the original distribution of inter-spindle intervals; only the order of the intervals was randomized. For each of the 1000 permutations of randomized data, the method described above for calculating mean probabilities was repeated and a grand mean for each plot was calculated.

To determine whether the probability of a sleep spindle was significantly greater in the immediate vicinity of a twitch, we calculated the peak probability of a sleep spindle within +3 s of an arm or leg twitch for the actual (observed) and shuffled (expected) data. Paired t tests were conducted on the 10 limb-age combinations; alpha was set at .05 and a Bonferroni procedure was used to correct for multiple comparisons.

## DATA AND CODE AVAILABILITY

Coded data and a subset of videos will be shared on Databrary (https://nyu.databrary.org). Custom MATLAB scripts used in this study will be shared on Github (https://github.com).

## Supplemental Information

**Figure S1.**
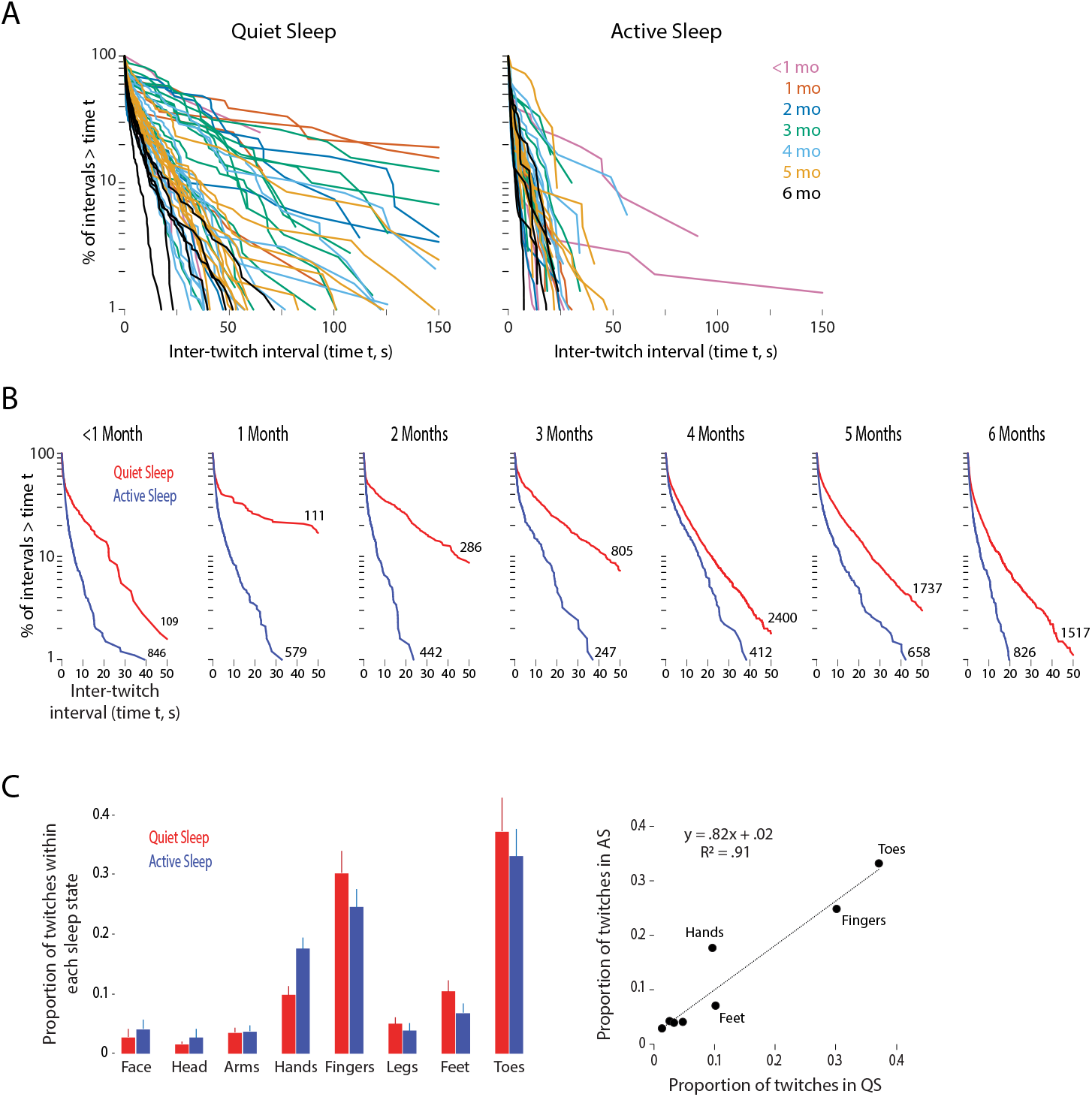
Spatiotemporal distribution of twitches during QS and AS. (A) Log-survivor plots comparing inter-twitch intervals (ITIs) between periods of QS (left) and AS (right) for every individual sleep session. Plots are color-coded by the age of the infant during each session. The sample sizes for each individual plot range from 4 to 657 (median=72.5) for QS and 6 to 463 (median=66) for AS. (B) Log-survivor plots to compare inter-twitch intervals (ITIs) between periods of QS (red lines) and AS (blue lines) within each age group. ITIs were pooled across sleep sessions within each age. Numbers adjacent to each plot indicate the number of ITIs. All within-age pairwise comparisons are significant using the Mantel-Cox log-rank test (X^2^s >19.7, ps<.00715 after Bonferroni correction). (C) Distribution of QS- and AS-related twitches across the body in 5-month-old infants. Left: For QS (red) and AS (blue), the mean proportions of twitches that occurred in the face, head, arms (shoulders/elbows), hands, fingers, legs (hips/knees), feet, and toes are shown. Data are from eight infants who had bouts of both QS and AS during the sleep session. Right: Scatterplot showing the proportion of twitches in QS and AS for each body segment for the data in the left panel. The best-fit line, regression equation, and R^2^ value are also shown.

**Figure S2.**
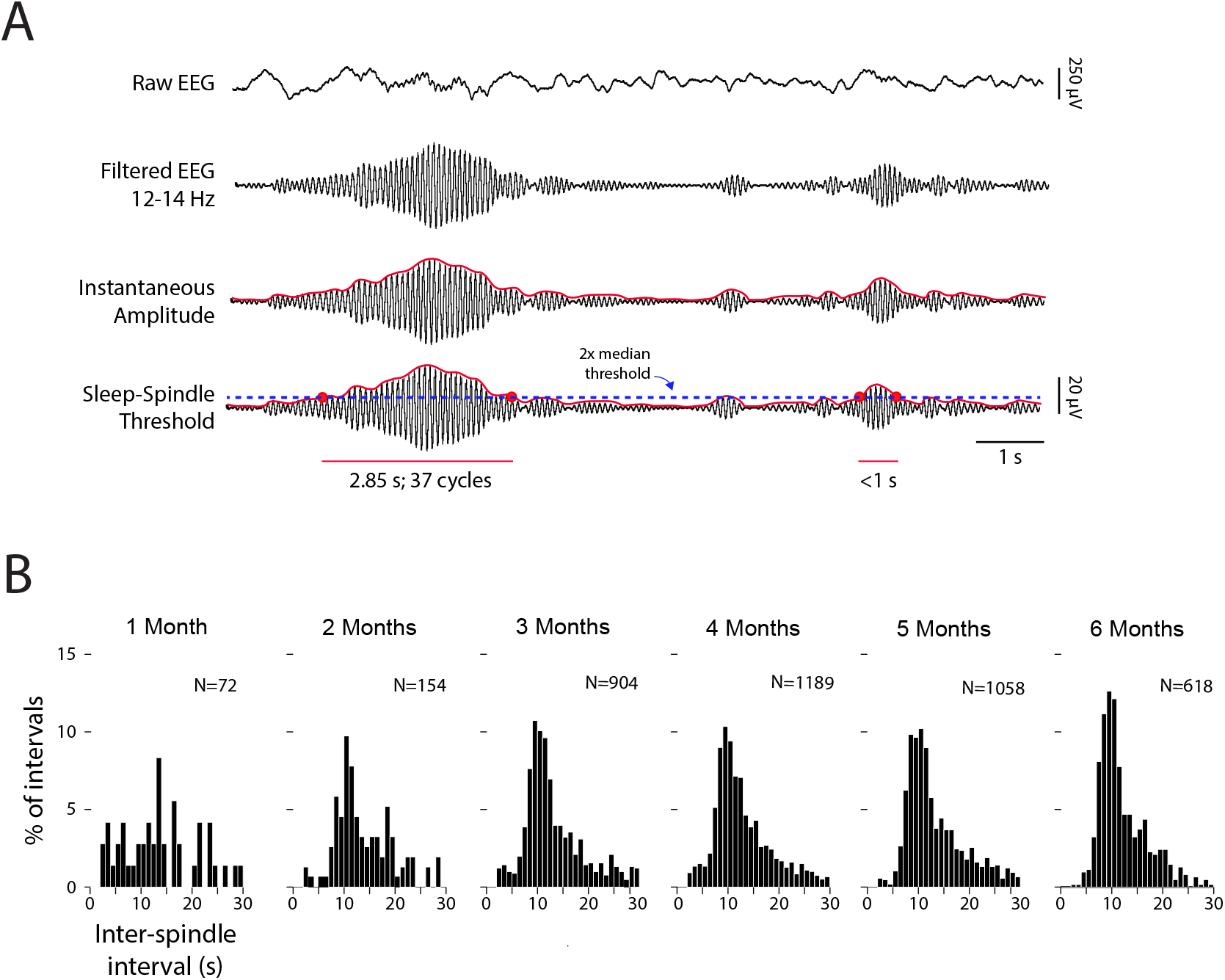
Detection and temporal distribution of sleep spindles. (A) Starting with the raw EEG signal (top row), sleep spindles were detected by first band-pass filtering the signal at 12-14 Hz (second row). Using a Hilbert transform of the filtered data, a continuous record of spindle amplitude was generated (third row). Sleep spindles were identified when signal amplitude exceeded a threshold of 2x the median amplitude for at least 1 s (fourth row). (B) Frequency distributions of inter-spindle intervals during QS within each age group. Intervals were pooled across sleep sessions within each group. The number (N) of sleep spindles contributing to each plot is also shown.

**Figure S3.**
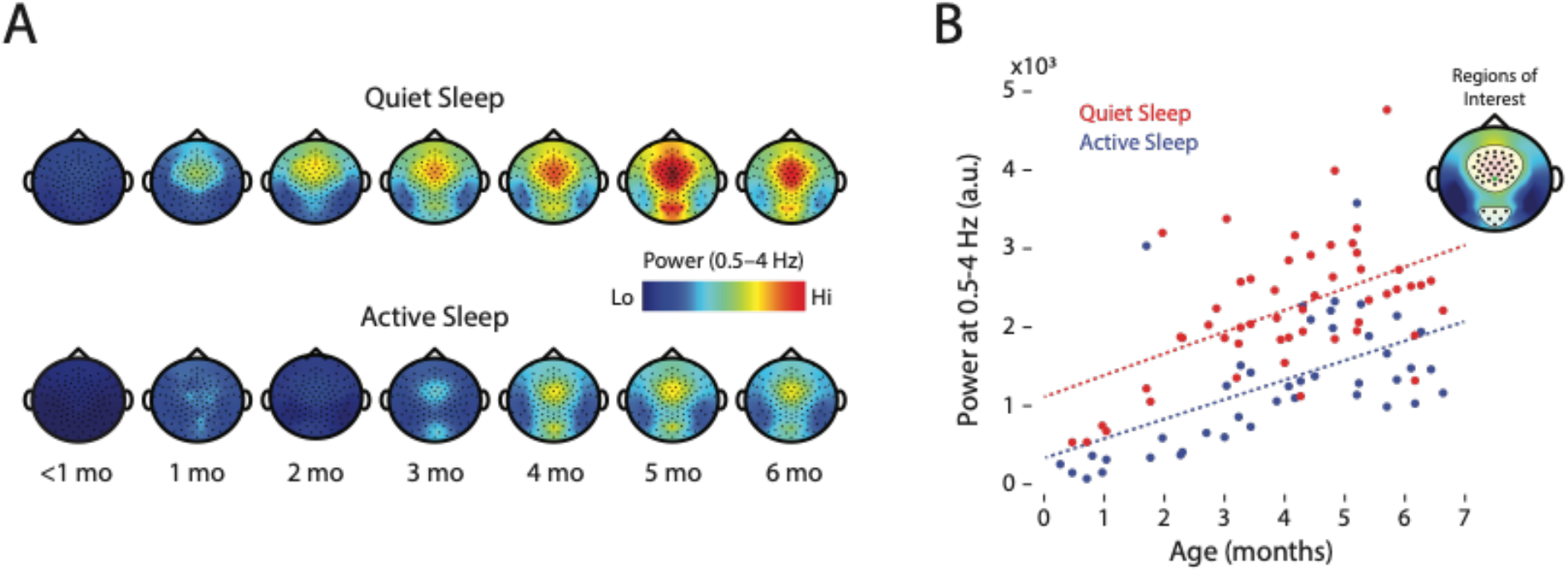
State-dependent increase in delta power over development. (A) Topoplots showing power in the delta range (0.5-4 Hz) during QS and AS. Data averaged across sleep sessions at each age. (B) Power at delta frequency (0.5-4 Hz; arbitrary units) averaged within a region of interest (inset; Cz electrode indicated by a green dot) during QS (red) and AS (blue) for each sleep session. The dotted lines show the model-predicted estimates for population means by age and sleep state. This model revealed significant main effects of sleep state (B=-895.44, SE=141.79, t(71.0)=-6.32, p<.0001) and age (B=264.93, SE=43.43, t(63.1)=6.10, p<.0001), but not a significant interaction (B=-28.12, SE=83.33, t(73.0)=-0.34, p=.74).

**Figure S4.**
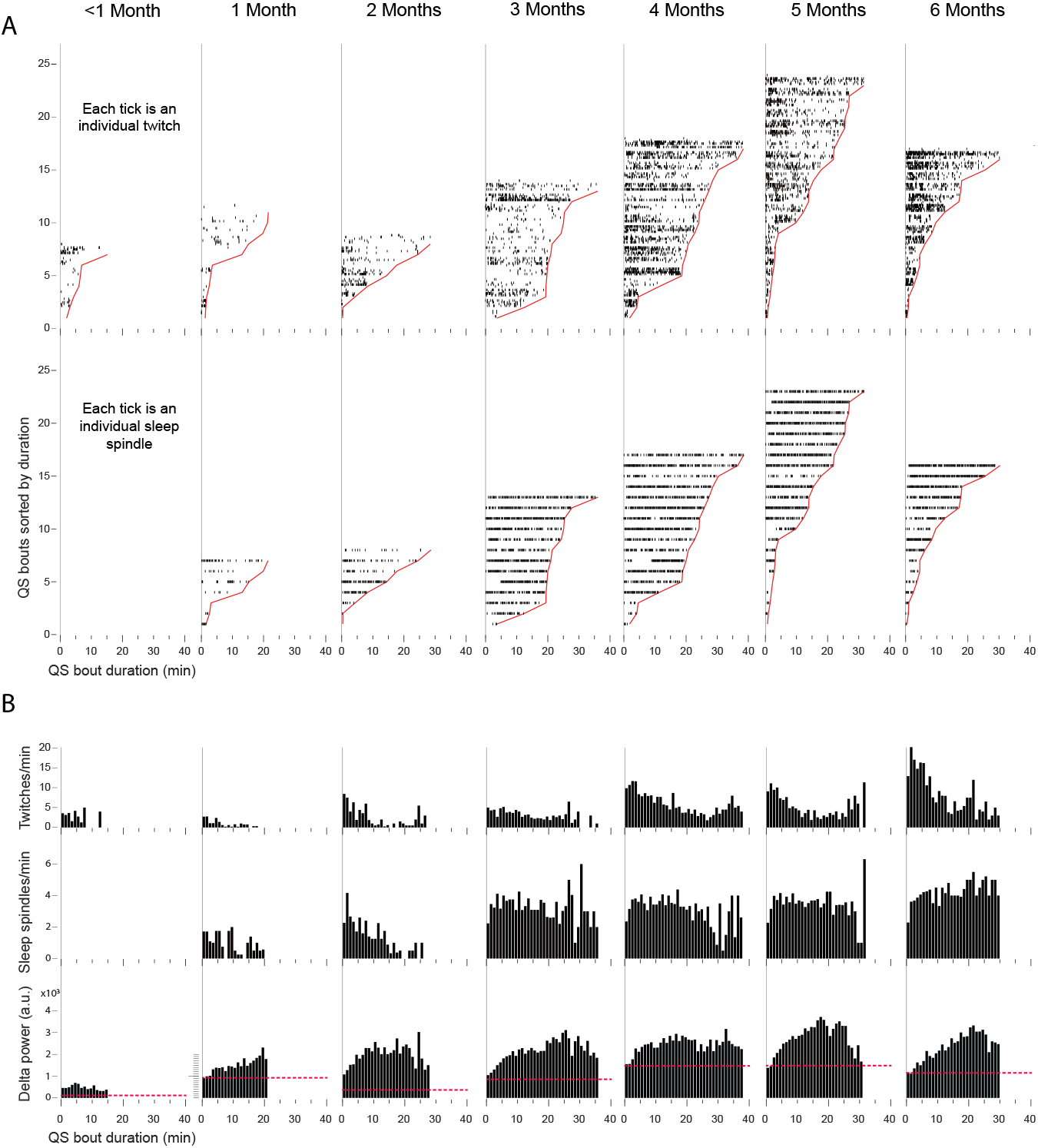
Temporal distribution of twitches, sleep spindles, and delta power across individual bouts of QS. (A) For each age group, QS bout duration is shown on the x-axis and QS bout number is shown on the y-axis, with bouts sorted from shortest to longest. The solid red lines show the duration of each bout. The black ticks show the occurrence of every twitch (top row) and sleep spindle (bottom row; detected at a single electrode site in somatosensory cortex) within the bout. For twitches, individual rows are devoted to twitches of the face, head, left and right arms, and left and right legs. (B) Same as in A except that the data are converted to rates (events/min) for the twitches (top row) and sleep spindles (middle row) for each 1-min bin. In the bottom row, mean delta power for each 1-min bin of the QS bout is also shown. For comparison, median delta power during AS for each age group is shown as a red dashed horizontal line.

**Table S1.** Summary information for the infant subjects and basic sleep data within each age group. Unless otherwise indicated, data are presented as mean ± sem; ranges are presented in parentheses. * Medians are presented for data that were not normally distributed (Shapiro-Wilk test, p<.05)

|  | <1 month | 1 month | 2 months | 3 months | 4 months | 5 months | 6 months |
| --- | --- | --- | --- | --- | --- | --- | --- |
| No. of sleep sessions<br>(M=male; F=female) | 5<br>(3M, 2F) | 3<br>(2M, 1F) | 5<br>(4M, 1F) | 12<br>(10M, 2F) | 12<br>(9M, 3F) | 10<br>(9M, 1F) | 6<br>(6M, 0F) |
| Postpartum Age | 0.63 ± 0.12<br>(0.25-0.95) | 1.60 ± 0.20<br>(1.03-1.94) | 2.57 ± 0.12<br>(2.26-2.87) | 3.45 ± 0.10<br>(3-3.97) | 4.44 ± 0.08<br>(4.06-4.81) | 5.43 ± 0.09<br>(5.14-5.87) | 6.29 ± 0.08<br>(6.1-6.61) |
| Weight (kg) | 4.07 ± 0.31<br>(3.35-5.17) | 5.21 ± 0.51<br>(4.05-6.10) | 5.54 ± 0.36<br>(4.26-6.30) | 6.43 ± 0.24<br>(4.99-7.96) | 7.07 ± 0.23<br>(5.45-8.41) | 7.90 ± 0.34<br>(5.53-9.92) | 7.68 ± 0.33<br>(6.18-8.5) |
| Height (cm) | 54 ± 1.67<br>(48-58) | 58 ± 1.63<br>(54-62) | 62.3 ± 2.29<br>(57-68.5) | 63.75 ± 1.17<br>(55.6-71) | 66.54 ± 1.26<br>(62-76) | 68.82 ± 1.03<br>(65-75) | 66.42 ± 0.66<br>(64-68.5) |
| Session Length (min) | 48.87 ± 10.52<br>(37.56-69.89) | 54.49 ± 6.44<br>(37.97-68.09) | 42.82 ± 6.8<br>(27.74-57.6) | 32.83 ± 2.71<br>(20.91-53.18) | 44 ±<br>4.22<br>(21.12-79.57) | 43.71 ± 5.87<br>(19-55.16) | 36.79 ± 7.14<br>(20.03-66.78) |
| AS Total Duration (min) | 12.2 ± 2.67<br>(5.61-20.52) | 12.47 ± 6.30<br>(5.03-31.27) | 5.36 ± 2.68<br>(0.0-14.58) | 2.72 ± 0.93<br>(0.0-9.54) | 3.69 ± 1.25<br>(0.0-14.49) | 5.3 ± 1.64<br>(0.0-14.42) | 4.45 ± 1.17<br>(0.0-7.76) |
| QS Total Duration (min) | 8.79 ± 3.85<br>(0-18.82) | 26.04 ± 2.00<br>(19.58-32.78) | 19.63 ± 5.30<br>(0-28.99) | 22.94 ± 1.78<br>(12.41-36.07) | 29.80 ± 2.96<br>(18.84-56.98) | 28.08 ± 2.55<br>(13.93-42.71) | 27.17 ± 6.79<br>(12.54-56.91) |
| Twitches/Min During AS | 14.93 ± 4.14<br>(2.07-24.43) | 13.02 ± 1.70<br>(9.14-17.23) | 18.05 ± 6.89<br>(10.7-31.82) | 10.42 ± 2.38<br>(3.05 – 21.4) | 11.35 ± 1.74<br>(3.77-18.75) | 12.33 ± 1.95<br>(2.09-19.33) | 20.76 ± 5.31<br>(9.41-37.7) |
| Twitches/Min During QS | 2.93 ± 1.40<br>(0.37-5.21) | 1.32 ± 0.82<br>(0.36-3.78) | 3.06 ± 1.12<br>(1.19-6.17) | 3.3 ± 0.77<br>(0.84-9.37) | 6.59 ± 0.93<br>(2.02-10.79) | 5.66 ± 0.44<br>(2.16 – 7.12) | 11.72 ± 2.15<br>(7.28-21.77) |
| Inter-Twitch Interval AS (s)* | 1.14 | 1.12 | 0.92 | 1.19 | 1.44 | 1.18 | 0.85 |
| Inter-Twitch Interval QS (s)* | 1.59 | 2.38 | 1.10 | 6.71 | 2.47 | 3.08 | 1.288 |
| Sleep Spindles/Min | 0 | 0.71 ± 0.37<br>(0-1.75) | 1.32 ± 0.50<br>(0.59-2.79) | 2.86 ± 0.20<br>(1.91-4.38) | 3.22 ± 0.27<br>(1.67-4.41) | 3.24 ± 0.29<br>(1.53-4.53) | 3.27 ± 0.43<br>(1.64-4.09) |
| Sleep Spindle Duration (s) | NA | 1.79 ± 0.28<br>(1.35-2.33) | 2.15 ± 0.38<br>(1.41-3.15) | 2.36 ± 0.15<br>(1.56-3.29) | 2.48 ± 0.11<br>(1.68-2.93) | 2.48 ± 0.14<br>(1.61-3.3) | 2.47 ± 0.16<br>(2.02-3.18) |
| Number of Cycles | NA | 23.57 ± 3.36<br>(18.41-29.88) | 28.12 ± 5.23<br>(18.72-42.36) | 31.16 ± 2.07<br>(20.72-45.09) | 32.60 ± 1.53<br>(22.51-39.48) | 32.73 ± 1.83<br>(22.1-43) | 32.47 ± 2.07<br>(27.1-41.65) |
| Inter-Spindle Interval (s)* | 0 | 17.99 | 16.36 | 12.24 | 12.21 | 12.01 | 11.08 |

**Movie S1**. Video showing twitching during AS (left) and QS (right) in a 6-month-old human infant. Most of the twitches occur in the hands, fingers, feet, and toes. Each recording is 30 s in duration and playback speed is at 3x. Two sleep spindles are shown during QS, recorded at electrode C3 and bandpass filtered (12-14 Hz).

